# Indium Tin Oxide (ITO) Substrates Enable Coating-Free SEM Imaging and Simplified Preparation of Purified Fibrinogen Clots

**DOI:** 10.64898/2026.06.24.733738

**Authors:** Can Cai, Caela Flake, Arezoo Nameny, Nathan E. Hudson, Brittany E. Bannish, Martin Guthold

## Abstract

**Background:** Scanning electron microscopy (SEM) is widely used to determine fibrin fiber structural properties such as fiber diameter and fiber length. However, conventional SEM preparation protocols are time-consuming and typically require conductive sputter coating. The coating process introduces an additional layer onto the sample surface and may influence measurements of nanoscale fiber structure. Furthermore, preparation of purified fibrinogen clots often follows protocols originally developed for plasma clots, resulting in unnecessary processing steps.

**Objective:** To evaluate indium tin oxide (ITO) as a flat, conductive substrate for SEM imaging of fibrin fibers, investigate the effects of sputter coating on measured fiber diameter, and develop a simplified SEM preparation protocol for purified fibrinogen clots.

**Methods:** Platelet-poor plasma clots and purified fibrinogen clots were formed on ITO substrates and imaged by SEM following 0 s, 45 s, or 90 s sputter coating. Fibrin fiber diameters were quantified and compared across coating conditions. For purified fibrinogen clots, an ITO-based simplified preparation protocol, in which clots were formed and imaged directly on the conductive ITO surface, was compared with a previously developed, standardized SEM protocol, in which clots were formed in microtube lids and subsequently transferred onto carbon tape for imaging.

**Results:** Fiber diameter measurements were affected by sputter coating duration, with increasing coating time resulting in larger apparent fiber diameters. Plasma and purified fibrinogen clots exhibited distinct fiber diameter distributions and coating responses. For purified fibrinogen clots, the simplified ITO-based protocol produced fiber diameter measurements that were not significantly different from those obtained using the standardized lid-to-carbon-tape workflow when identical coating times were applied.

**Conclusions:** ITO provides a practical conductive substrate for SEM imaging of fibrin fibers and enables substantial simplification of purified fibrinogen clot preparation. When coating conditions are matched, the simplified ITO-based protocol yields fiber diameter measurements comparable to those obtained using the previously standardized lid-to-carbon-tape workflow. These findings support the use of ITO as an alternative conductive imaging substrate and provide a simplified workflow for SEM analysis of purified fibrinogen clots. By reducing washing and transfer steps, this workflow may also provide a useful platform for future controlled studies of fibrin interactions with added proteins or other associated components.

## Introduction

Blood clot formation is essential for hemostasis; however, abnormal clot formation and impaired clot breakdown are associated with numerous diseases, including myocardial infarction, ischemic stroke, venous thromboembolism, pulmonary embolism, diabetes, and cardiovascular disease (*1, 2*). Cardiovascular diseases remain the leading cause of death worldwide and account for approximately one-third of all global deaths. Because fibrin forms the primary structural scaffold of blood clots, understanding how fibrin structure changes under different physiological and pathological conditions is important for understanding clot function and disease progression (*3*).

Over the past several decades, electron microscopy has become one of the most important tools for studying blood clot ultrastructure. It has been used to investigate fibrin polymerization, protein–protein interactions, clot architecture, and structure–function relationships within fibrin networks (*4-6*). Numerous studies have shown that fibrin morphology is associated with disease state, clot mechanical properties, permeability, and susceptibility to fibrinolysis. Structural characteristics such as fiber diameter, fiber length, branching, network density, and pore size have all been reported to differ between healthy and disease-associated clots (*1-3, 7*).

Among the available imaging techniques, scanning electron microscopy (SEM) remains one of the most widely used methods for quantitative analysis of fibrin structure because it provides sufficient resolution to visualize individual fibrin fibers. SEM has been extensively applied to whole blood clots (*1-3, 7*)(*8*), platelet-poor plasma clots, and purified fibrinogen clots to quantify fibrin fiber diameter, fiber length, branching, and other structural parameters. In particular, fibrin fiber diameter is one of the most frequently reported measurements and is commonly used to compare clot structure across disease states, experimental conditions, and biochemical compositions (*9-12*).

Despite its widespread use, conventional SEM preparation of fibrin clots is labor-intensive and time-consuming. Typical protocols involve fixation, washing, dehydration, drying, mounting, conductive sputter coating, and imaging. Conductive coating is commonly required to prevent charging of biological samples during SEM imaging; however, the deposited metal layer introduces an additional nanoscale coating onto the fiber surface and may influence quantitative measurements of fiber diameter. This issue is particularly relevant when analyzing fibrin fibers, whose diameters often range from only tens to hundreds of nanometers (*6, 9, 13, 14*).

In addition, many SEM preparation procedures currently used for purified fibrinogen clots are adapted from protocols originally developed for plasma clots (*13, 15*). Because purified fibrinogen systems lack many plasma components, some of these preparation steps may be unnecessary and may increase preparation time without improving measurement quality.

Therefore, there is a need for simplified SEM workflows that reduce sample processing while maintaining reliable quantitative measurements.

Indium tin oxide (ITO) is an inexpensive, flat, conductive substrate that may provide an alternative approach for SEM imaging by reducing or eliminating the need for conductive sputter coating (*16*). In this study, we evaluate ITO substrates for SEM imaging of fibrin fibers, investigate the effects of sputter coating duration on measured fibrin fiber diameter, and develop a simplified preparation protocol for purified fibrinogen clots. We further compare this simplified ITO-based workflow with a previously standardized SEM protocol to determine whether reliable quantitative measurements can be obtained with fewer preparation steps.

## Materials and Methods

### Reagents

Platelet-poor, pooled normal human plasma was purchased from George King Bio-Medical (Product #0010). Peak 1 purified human fibrinogen was purchased from Enzyme Research Laboratories (Cat# P1 FIB). Human α-thrombin was obtained from Enzyme Research Laboratories (Cat# HT1002a; lot-specific activity: 3044 NIH U/mg). Plasma, fibrinogen, and thrombin stock solutions were divided into single-use aliquots and stored at −80 °C. Before each experiment, aliquots were thawed at 37 °C and used immediately.

### Clot formation surfaces and SEM imaging substrates

ITO-coated glass was used as a conductive substrate for both clot formation and SEM imaging. ITO substrates were unpatterned ITO-coated glass microscope slides (25 × 75 mm; Ossila, Sheffield, UK), which were scored and cut with a diamond scribe into six individual pieces before use. In the standardized lid-to-carbon-tape method, clots were formed in microtube lids following the previously described SEM preparation protocol (*9*), then removed from the lids and mounted onto carbon tape for SEM imaging.

### Fibrin clot formation

Plasma and purified fibrinogen clots were prepared at final fibrinogen and thrombin concentrations of 2.9 mg/mL and 1.0 NIH U/mL, respectively. Clot formation was initiated by combining the fibrinogen-containing solution with thrombin and CaCl_2_ in Tris-buffered saline (50 mM Tris, 100 mM NaCl, pH 7.4). Plasma clots were prepared with 20 mM CaCl_2_ to recalcify the citrate-anticoagulated plasma. Purified fibrinogen clots were prepared with 9.5 mM CaCl_2_ because no citrate was present in the purified system. All clots were polymerized for 2 h at room temperature before fixation.

### Experimental Design

The study consisted of four experimental comparisons designed to evaluate sputter-coating effects, the effect of using ITO as a conductive imaging surface, and the feasibility of simplifying the purified fibrinogen clot preparation workflow. An overview of the experimental design is provided in Table 1.

**Table 1.** Experimental design and study comparisons.

| ***Comparison*** | ***Sample*** | ***Workflow*** | ***Coating*** | ***Purpose*** |
| --- | --- | --- | --- | --- |
| *Coating time on ITO* | Purified fibrinogen | ITO + Simplified protocol | 0, 45, 90 s | Test simplified ITO method with different coating times |
| *Coating time on ITO* | Plasma | ITO + Standard wash/dehydration steps | 0, 45, 90 s | Test coating effect on ITO |
| *Protocol comparison* | Purified fibrinogen | ITO + Simplified protocol vs Standardized lid-to-carbon-tape | Matched | Test whether simplified ITO protocol gives comparable fiber diameters |
| *ITO vs standardized lid-to-carbon-tape* | Plasma | ITO + Standard wash/dehydration steps vs Standardized lid-to-carbon-tape | Matched | Test whether ITO changes measured fiber diameter |

### ITO-based simplified preparation of purified fibrinogen clots

For the simplified ITO protocol, 10 µL of purified fibrinogen clot reaction mixture was pipetted directly onto the ITO-coated surface of an indium tin oxide (ITO)-coated glass substrate placed in a 12-well plate. After polymerization for 2 h at room temperature, the clot was fixed on the substrate by complete immersion in 2 mL of 2% glutaraldehyde prepared in sodium cacodylate buffer (50 mM sodium cacodylate, 100 mM NaCl, pH 7.4) overnight. Samples were covered during polymerization and fixation to minimize evaporation. After fixation, the fixation buffer was carefully removed by pipette, and 2 mL of 100% ethanol was added to cover the clot on the ITO substrate. The ITO samples were then transferred with tweezers to a Tousimis Samdri-PVT-3D critical point dryer for critical point drying.

This simplified workflow (Fig. 1) was designed to reduce sample handling by omitting the repeated buffer washing steps and the graded ethanol dehydration series used in the previously standardized fibrin SEM preparation protocol (*9*).

**Fig. 1.**
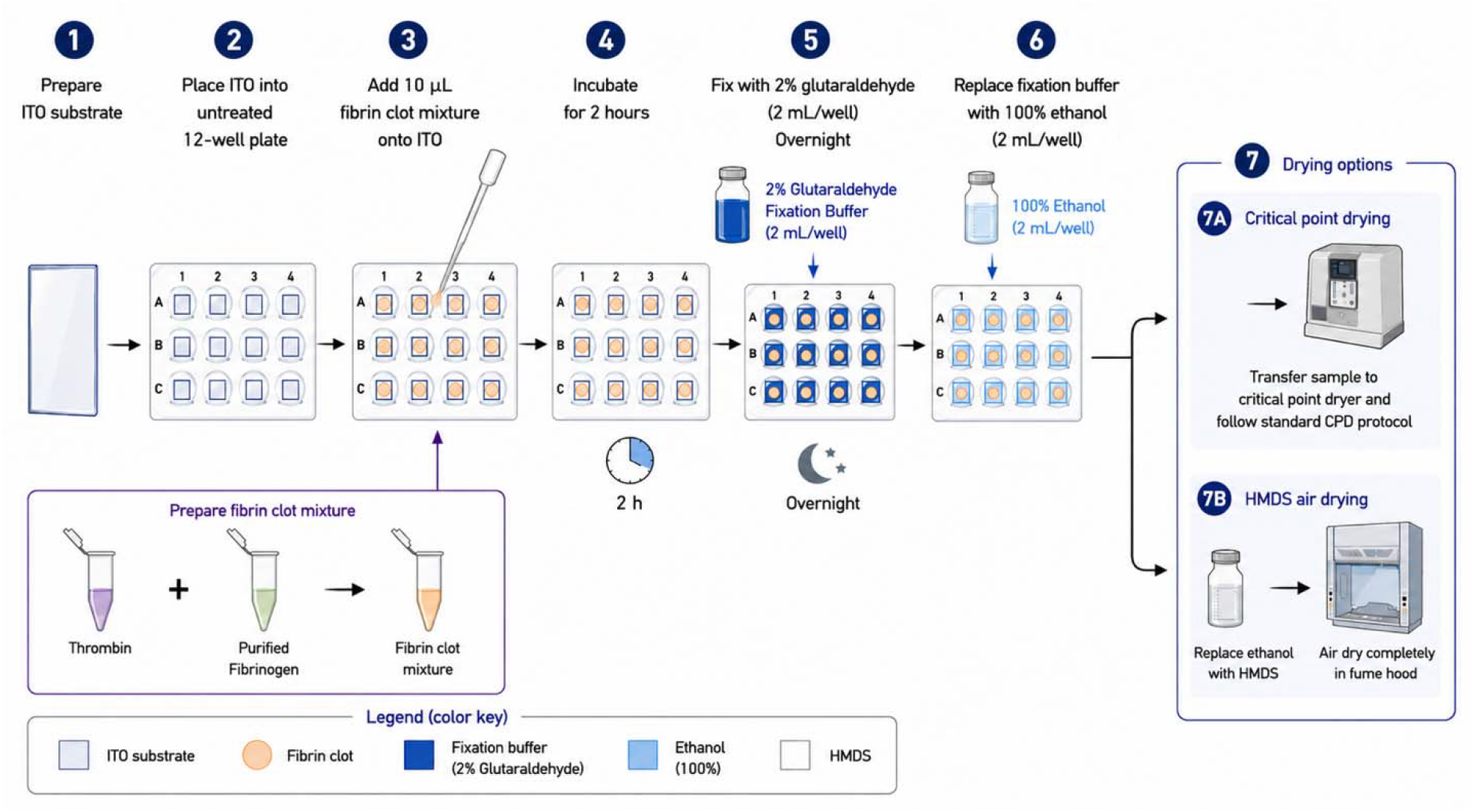
Simplified purified fibrinogen sample preparation for SEM. (This schematic was created with assistance from ChatGPT)

### Preparation of plasma clots on ITO

Plasma clots were prepared directly on ITO substrates to evaluate whether conductive ITO could support SEM imaging of plasma-derived fibrin networks with reduced or no sputter coating.

Because plasma clots contain additional plasma proteins, we included wash steps to remove non-fibrin components and leave primarily the fibrin fiber network. Plasma samples were therefore processed using a similar, previously described conservative standardized preparation workflow, except that clot formation was performed on ITO instead of in the microtube lid (*9*). After polymerization, samples were rinsed with sodium cacodylate buffer, fixed in 2% glutaraldehyde, rinsed again, dehydrated through increasing ethanol concentrations, and dried by critical point drying. Dried samples were either left uncoated or sputter-coated with gold for 45 s or 90 s.

### Comparison with the standardized lid-to-carbon-tape protocol

To determine whether the simplified ITO method produced fiber diameter measurements comparable to the previously standardized SEM method, purified clots were prepared using two workflows. In the first workflow, clots were formed directly on ITO substrates and processed using the simplified ITO method described above. In the second workflow, clots were formed in microtube lids (standardized lid-to-carbon-tape) and processed using the previously standardized wash, fixation, dehydration, drying, and sputter-coating procedure. For direct comparison, ITO and standardized lid-to-carbon-tape samples were sputter-coated for the same duration before SEM imaging. This design allowed the effect of the simplified ITO-based preparation to be evaluated without introducing differences in coating time.

### Sputter coating conditions

Dried samples were imaged either without sputter coating or after gold sputter coating using a Cressington Model 108 sputter coater. For coating-time experiments, samples were assigned to 0 s, 45 s, or 90 s coating conditions. For direct comparisons between the simplified ITO method and the standardized lid-to-carbon-tape method, both sample groups were coated for the same duration.

### SEM imaging

Samples were imaged using a Zeiss Gemini 300 scanning electron microscope under high vacuum at 30,000× magnification. For each experimental condition, two independently prepared clots were imaged. Eight non-overlapping fields of view were collected from each clot, yielding 16 SEM images per condition. Images were acquired from widely separated regions of the clot while avoiding the film-like edge region and obvious drying or handling artifacts.

For uncoated samples prepared on ITO substrates, imaging was performed at low accelerating voltage (≤5 kV) using the InLens secondary electron detector to minimize charging and enhance surface contrast. To further reduce charging and beam-induced artifacts, prolonged scanning of the same area was avoided (typically < 1 min per field of view), and regions showing obvious charging artifacts were excluded from analysis.

### Fiber diameter analysis

Fibrin fiber diameters were measured from SEM micrographs using the same analysis procedure for all experimental groups. Only clearly resolved fibers from the visible upper layer of the network were measured to avoid ambiguity from overlapping fibers. For each image, the median fiber diameter was calculated, and image-level median values were used for statistical comparisons. These comparisons were designed to evaluate sputter-coating effects, the effect of using ITO as a conductive imaging surface, and the feasibility of simplifying the purified fibrinogen clot preparation workflow.

## Results

### Coating time on ITO (Purified fibrinogen; simplified ITO protocol)

Sputter coating time produced a monotonic increase in the measured fibrin fiber diameter for purified fibrinogen clots prepared on ITO using the simplified protocol (Fig. 2). Across 16 SEM images per condition, the median image-level diameter increased from 69.6 nm (0 s) to 74.3 nm (45 s) and 89.4 nm (90 s). The coating-time effect was significant by Kruskal–Wallis (H = 30.40, p < 0.0001; n = 16 images/group), and Dunn’s multiple-comparisons test showed 0 s vs 45 s, p = 0.58; 0 s vs 90 s, p < 0.0001; 45 s vs 90 s, p = 0.0002.

**Fig. 2.**
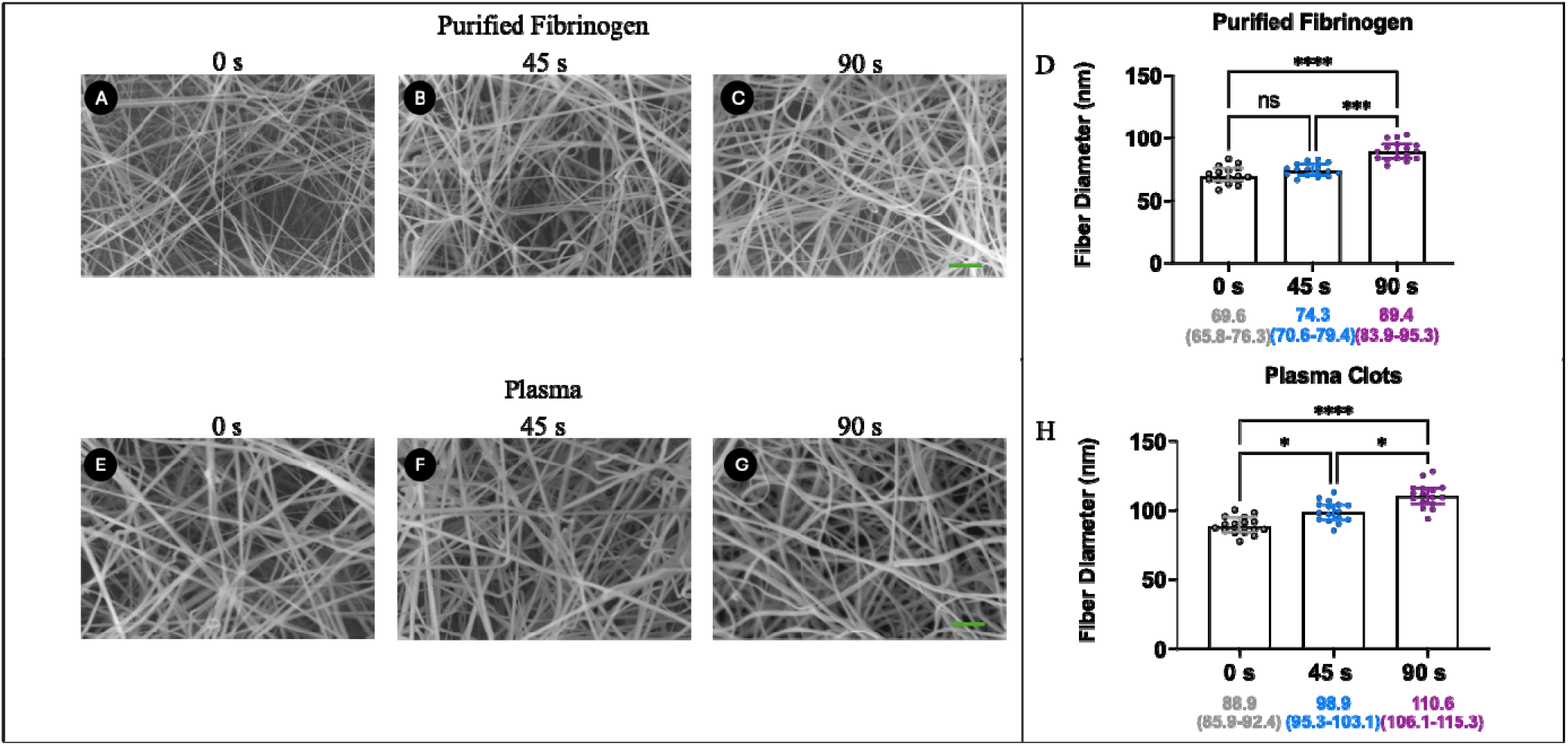
Coating-time dependence of fibrin fiber diameter measurements on ITO. (A–C) Purified fibrinogen clots (simplified ITO protocol, Fig. 1) with coating times of 0, 45, and 90 s. (D) Image-level medians of fiber diameter (n = 16 SEM images per timepoint); points are per-image medians; line = median across images; bars = 95% CI. (E–G) Plasma clots with coating times of 0, 45, and 90 s. (H) Image-level medians of fiber diameter (n = 16 per timepoint); points are per-image medians; line = median across images; bars = 95% CI.

### Coating time on ITO (Plasma; standard protocol on ITO)

Plasma clots prepared on ITO showed a similar coating-dependent increase in measured fiber diameter (Fig. 2). The median image-level diameter increased from 88.9 nm (0 s) to 98.9 nm (45 s) and 110.6 nm (90 s) across 16 SEM images per condition. Kruskal–Wallis indicated significant differences among coating times (H = 29.34, p < 0.0001; n = 16 images/group), and Dunn’s test yielded 0 s vs 45 s, p = 0.02; 0 s vs 90 s, p < 0.0001; 45 s vs 90 s, p = 0.02.

### Simplified ITO preparation vs standardized lid-to-carbon-tape (purified fibrinogen, 45 s coating)

The simplified ITO preparation for purified fibrinogen produced fiber diameter measurements comparable to the standardized lid-to-carbon-tape workflow (Fig. 3). Across 16 SEM images per condition (each point representing the median fiber diameter within one image), the median image-level diameter was 74.3 nm for ITO (45 s) and 74.9 nm for standardized lid-to-carbon-tape (45 s). A two-tailed Mann–Whitney test showed no significant difference between the two workflows (U = 116, p = 0.67).

**Fig. 3.**
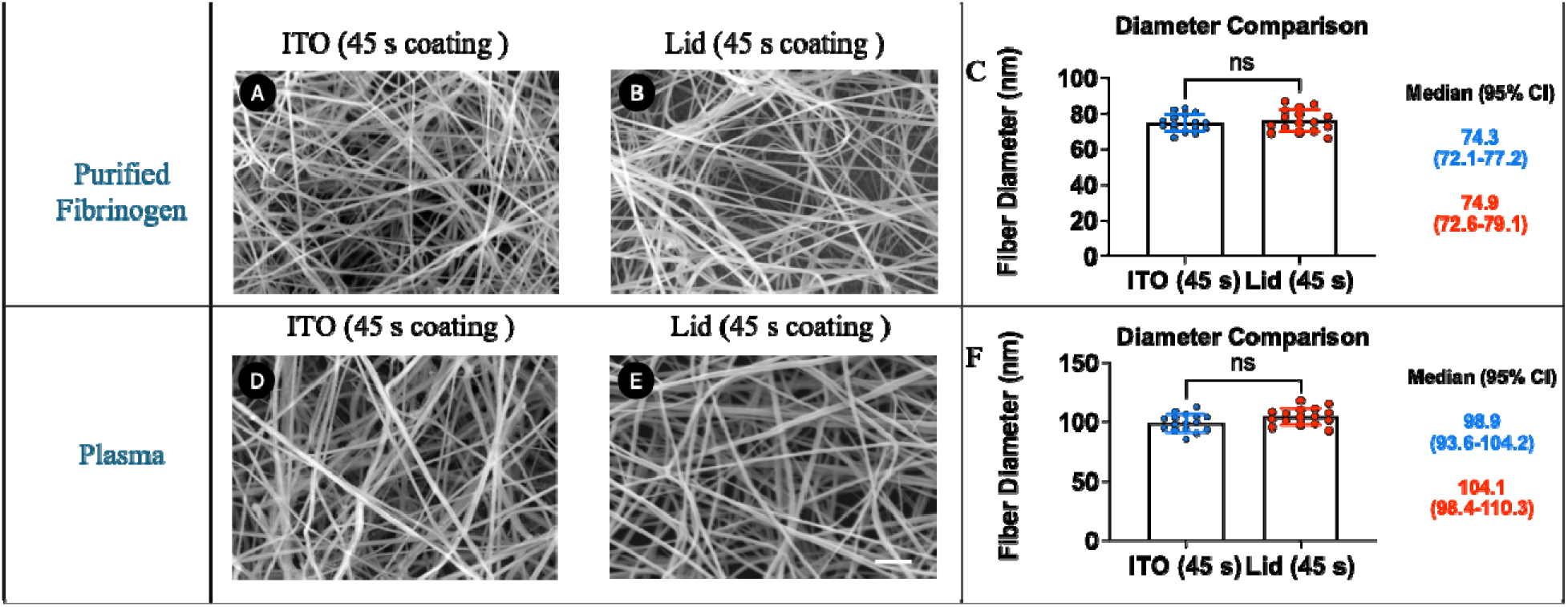
Validation of the simplified ITO protocol and comparison with the standardized lid- to-carbon-tape workflow. (A) Purified fibrinogen clot formed on ITO using the simplified protocol (45 s coating). (B) Purified fibrinogen clot formed in a microtube lid using the standardized lid-to-carbon-tape protocol (45 s coating). (C) Fiber diameter comparison for (A) vs (B): Each point is the median diameter from one SEM image (n = 16 images/condition); line = median across images; bars = 95% CI. (D) Plasma clot formed on ITO (45 s coating). (E) Plasma clot formed in a microtube lid (45 s coating). (F) Fiber diameter comparison for (D) vs (E): Each point is the median diameter from one SEM image (n = 16 images/condition); line = median across images; bars = 95% CI.

### ITO-based workflow vs standardized lid-to-carbon-tape workflow in plasma clots (45 s coating)

For plasma clots processed with the standardized workflow, measured fiber diameters on ITO were not significantly different from those formed on microtube lids (Fig. 3). The median image-level diameter across 16 SEM images per condition was 98.9 nm on ITO (45 s) and 104.1 nm on standardized lid-to-carbon-tape (45 s). This difference was not statistically significant by a two-tailed Mann–Whitney test (U = 79, p = 0.07).

## Discussion

This study addressed four practical questions relevant to reproducible SEM measurement of fibrin fiber diameter: (i) how sputter-coating time affects measured diameters on ITO, (ii) whether an ITO-based simplified workflow is feasible for purified fibrinogen clots, (iii) whether using ITO as the clot formation and imaging surface biases plasma fiber diameter measurements relative to the standardized lid-to-carbon-tape workflow, when the remaining preparation steps are held constant, and (iv) whether the simplified ITO workflow yields comparable results to the standardized lid-based workflow for purified fibrinogen.

Across both purified fibrinogen and plasma clots formed on ITO, sputter coating time produced a monotonic increase in the measured fiber diameter. For purified fibrinogen (simplified ITO protocol), the median of image-level medians increased from 69.6 nm (0 s) to 74.3 nm (45 s) and 89.4 nm (90 s), with significant differences driven primarily by the 90 s condition. Plasma clots formed on ITO showed a similar shift from 88.9 nm (0 s) to 98.9 nm (45 s) and 110.6 nm (90 s). These results reinforce that sputter coating is not a neutral imaging step: increasing coating time can introduce a systematic, directionally consistent bias in apparent fibrin fiber diameter. The magnitude of the shift is consistent with the expected addition of a nanoscale conductive gold layer during sputter coating; however, the exact coating thickness was not directly measured in this study and can vary with sputtering current, chamber pressure, working distance, and sample geometry. Thus, the observed diameter increases likely reflect, at least in part, the added coating layer rather than underlying structural changes in the fibrin network. Practically, this argues that coating conditions must be tightly standardized and explicitly reported whenever fiber diameter is used as a quantitative readout across experiments or laboratories.

A central motivation of using ITO is that its conductive surface can reduce charging artifacts and enable imaging with reduced or even no sputter coating, which would eliminate one major source of measurement variability. In our SEM workflow, uncoated imaging on ITO was performed at low accelerating voltage (≤5 kV) using the InLens detector, and with limited dwell time per field of view to minimize charging and beam-induced artifacts. However, conductivity of the substrate does not guarantee charge dissipation for all sample geometries. If portions of the clot are elevated or poorly attached relative to the conductive ITO surface, such as thick deposits, folded regions, or detached material, those regions may still charge during SEM imaging. We therefore used a small reaction volume (10 µL) to form shallow dome-shaped clots on the ITO surface and selected imaging regions that appeared well-coupled to the conductive substrate.

Assuming a sample area of 25 mm^2^, the average clot height is 40 µm.

The simplified ITO-based preparation for purified fibrinogen produced measurements comparable to the standardized lid-based workflow at matched coating time (45 s). The median image-level fiber diameter was 74.3 nm on ITO versus 74.9 nm on standardized lid-to-carbon-tape (n = 16 images per condition; Mann–Whitney p = 0.6689), indicating no detectable bias introduced by the simplified ITO-based workflow in the purified system. This supports the use of the simplified ITO protocol as a practical alternative for purified fibrinogen clots, reducing handling steps while preserving quantitative readouts.

In contrast, plasma clots contain additional plasma proteins and non-fibrin components that can remain associated with the network after polymerization. As a result, conservative wash steps remain important if the goal is to measure fibrin fiber diameters comparably to established standardized workflows. Consistent with this, we retained wash and graded dehydration steps for plasma in the standard protocol (and as illustrated in Supplementary Fig. 1), to omit the risk of leaving residual material that can obscure fibers or confound diameter measurement. Importantly, when plasma samples were processed with the same standardized workflow, the choice of ITO-based workflow vs standardized lid-to-carbon-tape workflow did not significantly alter measured diameters at 45 s coating (98.87 nm on ITO vs 104.1 nm on standardized lid-to-carbon-tape; n = 16 images per condition; Mann–Whitney p = 0.0671). This suggests that, under a controlled protocol, the use of ITO as the clot formation and imaging surface is unlikely to be a dominant driver of between-study variation in plasma fiber diameter, and that protocol differences (including coating conditions and preparation steps) are more consequential.

Although residual non-fibrin material is undesirable when the goal is standardized fibrin fiber diameter measurement, this observation may also be useful in a different context. The accumulation of material on or around fibrin fibers in minimally washed plasma samples suggests that reduced-washing workflows may help preserve weakly associated plasma components that are otherwise removed during conventional preparation. Thus, while conservative washing remains preferable for quantitative diameter measurements in plasma, simplified or minimally washed ITO-based preparation may provide a useful platform for studying interactions between fibrin fibers and associated proteins or other soluble components in targeted experiments. In purified fibrinogen systems, where fewer non-fibrin components are present, the simplified ITO workflow may be especially useful for controlled studies in which defined proteins are added to fibrinogen before polymerization. Because the clot can be formed and imaged directly on the conductive ITO surface with fewer washing and transfer steps, this approach may help preserve protein–fibrin associations and reduce preparation-induced loss or disruption of loosely associated material.

Together, these findings highlight two sources of variability that can plausibly contribute to the wide spread of “normal” fiber diameters reported across laboratories: differences in sputter-coating conditions and differences in preparation workflows (particularly for complex samples such as plasma). Using conductive ITO substrates can reduce charging constraints and enable protocol simplification in purified systems, but plasma samples still benefit from conservative washing and dehydration steps to isolate fibrin fibers for consistent quantification.

## Conclusion

Using ITO-coated glass as a conductive substrate enables a simplified SEM preparation workflow for purified fibrinogen clots without altering measured fiber diameter relative to the standardized lid-based method at matched coating time. For plasma clots, the ITO-based clot formation and imaging workflow did not significantly change measured fiber diameters compared with the standardized lid-to-carbon-tape workflow when the remaining preparation steps were matched. However, washing and graded dehydration steps remain important for plasma samples because additional plasma proteins may obscure fibrin fibers or confound diameter measurements.

Sputter-coating time systematically increases the measured fiber diameter on ITO for both purified fibrinogen and plasma, consistent with the expected deposition thickness per unit time. Therefore, coating conditions should be standardized and explicitly reported to support meaningful comparisons across experiments and laboratories. The simplified ITO workflow may also be useful for future controlled studies of fibrin interactions with defined added proteins or associated components, because reduced washing and transfer steps may help preserve loosely associated material. Finally, while ITO can mitigate charging and facilitate reduced-coating imaging at low accelerating voltage, reliable uncoated imaging depends on maintaining close coupling between the sample and the conductive surface, motivating thin deposits and careful selection of imaging regions

## Declaration of AI and AI-assisted technologies in the writing process

During the preparation of this work, the author(s) used ChatGPT (OpenAI) to improve English grammar and clarity and to assist with the creation of the schematic illustration shown in Figure

1. After using this tool, the author(s) reviewed and edited the content as needed and take full responsibility for the content of this manuscript.

## Author contributions

Conceptualization: CC, CF, AN, NEH, BEB, MG

Data curation: CC, CF, AN, NEH, MG

Formal analysis: CC, CF, AN, MG

Funding acquisition: NEH, BEB, MG

Investigation: CC, CF, AN, NEH, BEB, MG

Methodology: CC, CF, AN, NEH, BEB, MG

Project administration: NEH, BEB, MG

Resources: NEH, BEB, MG

Software: CC Supervision: MG

Validation: CC, CF, AN, NEH, BEB, MG

Visualization: CC

Roles/Writing - original draft: CC, MG

and Writing - review & editing: CC, CF, AN, NEH, BEB, MG

## Acknowledgements

This work was supported by National Institutes of Health grants 2R15HL148842-02. The content is solely the responsibility of the authors and does not necessarily represent the official views of the National Institutes of Health. We thank Dundappa Mumbaraddi from the Department of Chemistry at Wake Forest University for his assistance with SEM imaging.

