## Supplement for "Indium Tin Oxide (ITO) Substrates Enable Coating-Free SEM Imaging and Simplified Preparation of Purified Fibrinogen Clots"

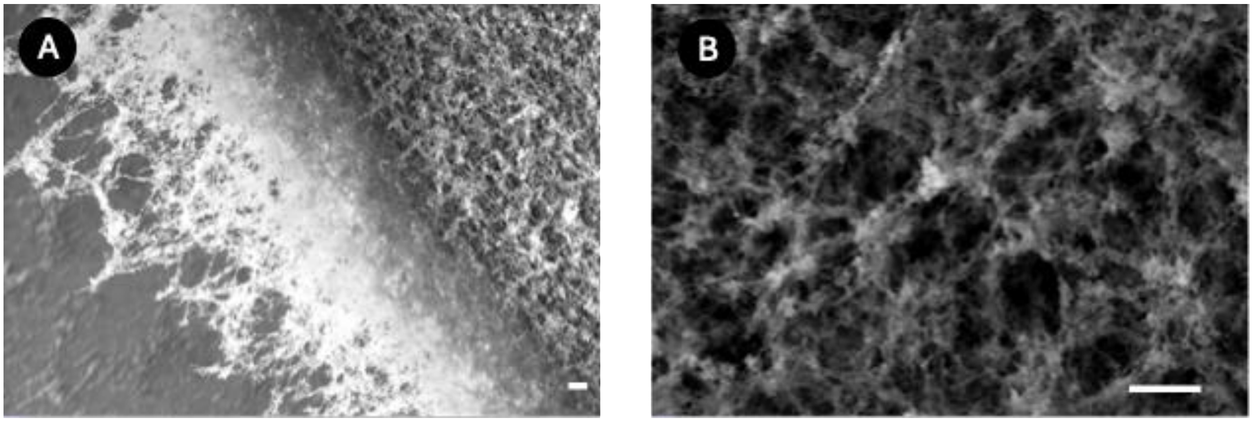


**Supplementary Figure 1. Plasma clot prepared using the simplified ITO workflow without washing and graded dehydration. Scale bar, 1 µm.** (A) Low-magnification SEM image showing the edge of a plasma clot prepared using the simplified ITO protocol. The sample contains abundant residual material associated with or deposited around the fibrin network, likely reflecting retained plasma components after omission of washing steps. (B) Higher-magnification SEM image showing that residual material obscures individual fibrin fibers, making the sample unsuitable for reliable fiber diameter quantification.

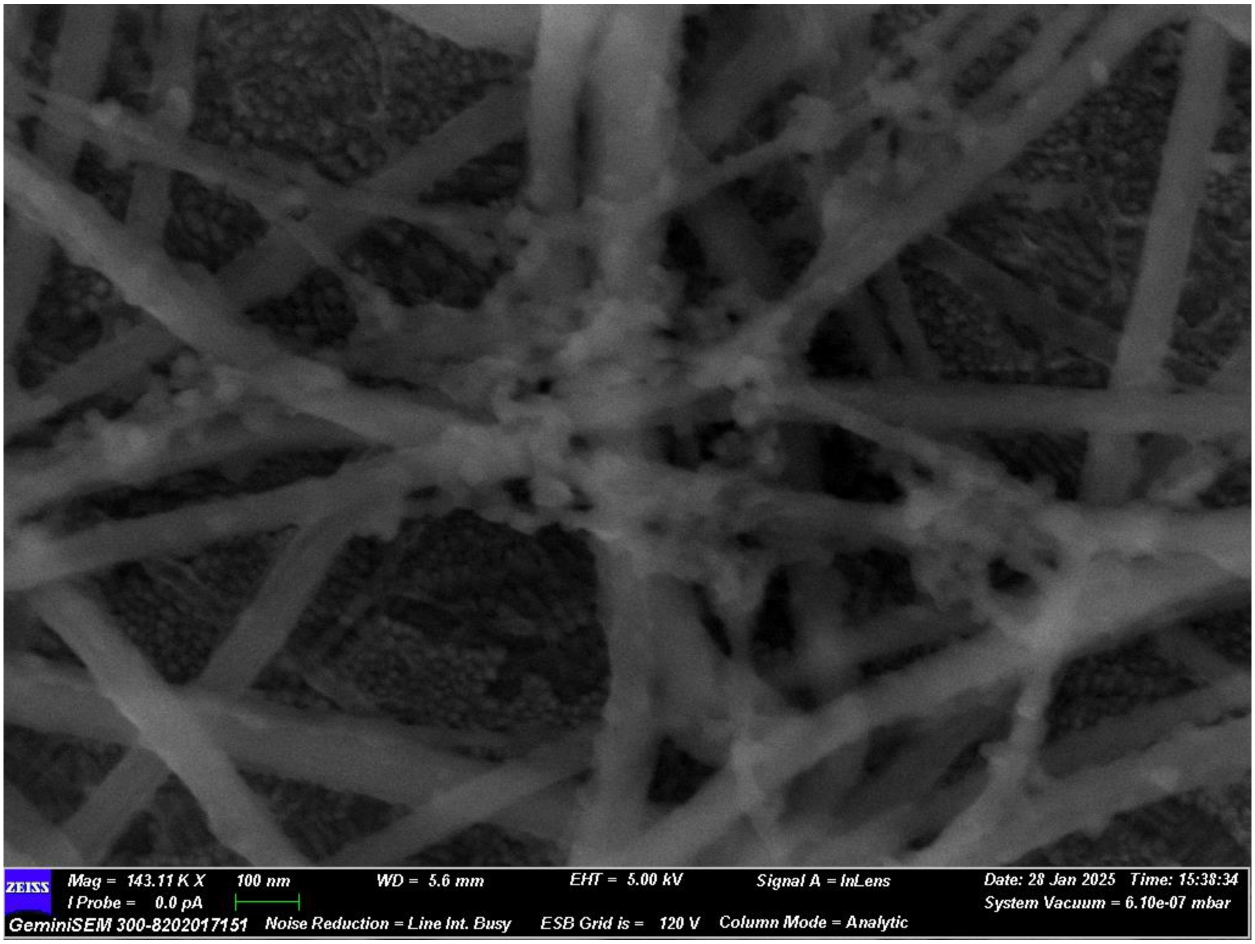


**Supplementary Figure 2. Zoomed-in image of plasma clot prepared using the simplified ITO workflow without washing and graded dehydration.** In a higher-resolution SEM image from a region with less residual material, individual fibrin fibers were more clearly visible, but additional material could still be observed attached to or deposited on the fiber surface.
